# Neurobehavioral impacts of the autism risk gene, WAC: Studies involving C. elegans and Mice

**DOI:** 10.64898/2026.03.02.709202

**Authors:** Napissara Boonpraman, Da-Woon Kim, Elena Tislerics, Janki Barot, Dariangelly Pacheco-Cruz, Nathan C Kuhn, Daniel Vogt, Shreesh Raj Sammi

## Abstract

Autism Spectrum Disorder (ASD) is a neurodevelopmental disorder characterized by a broad spectrum of behavioral impairments. While multiple genetic and environmental factors are attributed to its cause, biological underpinnings are still poorly understood. We investigated an ASD-associated gene, WAC, for its neurobehavioral aspects using *C. elegans* and mice. Studies of *C. elegans* with *wac* gene deletions (*wac-1.1* and *wac-1.2*) showed enhanced acetylcholine-associated behavior, as indicated by the aldicarb assay. No alteration in acetylcholine levels or acetylcholinesterase activity was observed. Upon further investigation, we found that the elevated cholinergic transmission resulted from increased activity of nicotinic acetylcholine receptors (nAChRs). Additionally, we observed reduced motility and dopamine-associated behaviors, along with a reduced ability to switch from crawling to swimming, a serotonin-dependent behavior. Upregulation in mRNA expression of the *lev-1* gene was observed. Conversely, a feedback-counterbalancing response in the form of downregulated genes, *acr-2, unc-17, unc-63*, and *unc-50*, was also observed. Surprisingly, *lev-1* RNAi did not reverse the enhanced cholinergic transmission in PHX2587 worms, indicating the involvement of other players. To validate our findings, we also assessed CHRNA7 levels in *Wac*^+/-^ mice. While some genetic compensation was observed in heterozygous mice, we found a direct, inverse correlation between *Wac* mRNA expression and CHRNA7 levels in the mouse brain cortex, corroborating our findings from *C. elegans*. Overall, these studies indicate that *wac* gene deletion in *C. elegans* exhibits a neurotransmitter alteration that is relatable to ASD.

**Graphical Abstract:** 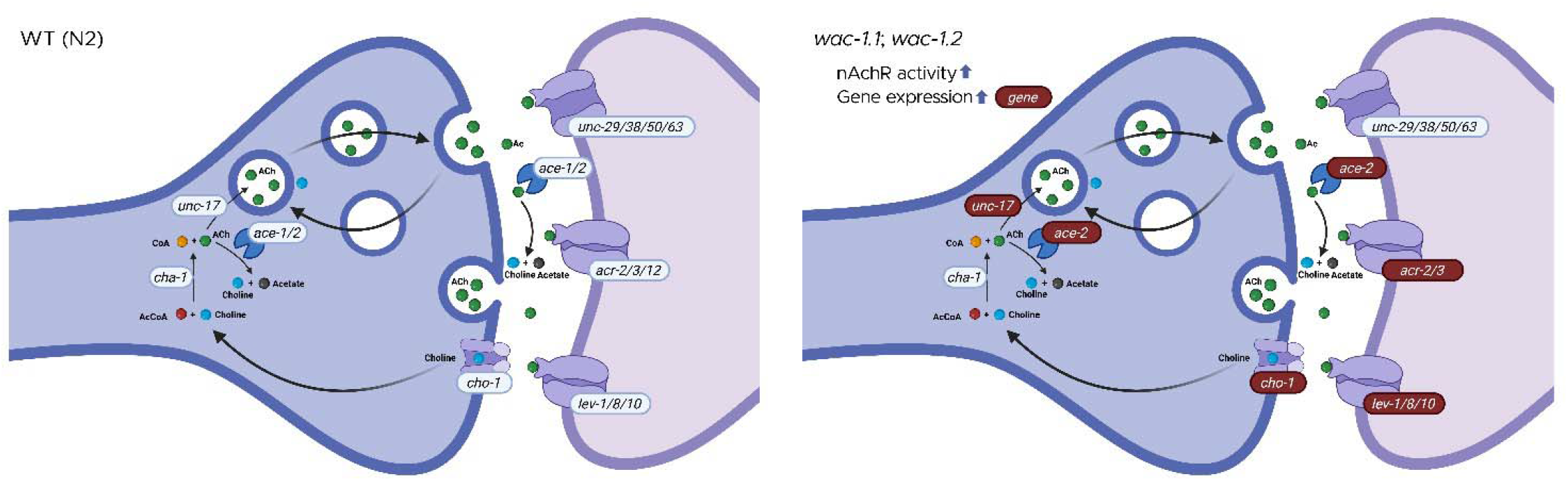

## Introduction

The WAC (WW domain–containing adaptor with coiled-coil) gene encodes a multifunctional protein characterized by WW and coiled-coil domains, structural motifs that commonly mediate protein–protein interactions[1]. WAC has been implicated in several fundamental cellular processes, including protein homeostasis[2,3], chromatin organization[4], transcriptional regulation[4,5] and alterations to behavior and cognition[6,7]. Despite these proposed roles, the precise biological functions of WAC and its impact on brain development and resulting behaviors remain obscure. Pathogenic variants in WAC cause DeSanto–Shinawi syndrome (DESSH), a rare neurodevelopmental disorder characterized by craniofacial dysmorphia, hypotonia, and global developmental delay. Individuals with DESSH frequently exhibit neuropsychiatric comorbidities, including attention deficit hyperactivity disorder (ADHD), autism-related traits, and increased seizure susceptibility, highlighting a critical role for WAC in brain development and function[7-9]. The phenotypic spectrum of DESSH suggests that WAC dysfunction disrupts neural pathways essential for cognition and behavior[7]. Consistent with this clinical presentation, WAC has also emerged as a high-confidence autism spectrum disorder (ASD) risk gene[9-14].

ASD is a neurodevelopmental condition defined by impairments in social communication and restricted and repetitive behaviors[15]. Large-scale genetic studies have identified de novo mutations in WAC in individuals with ASD and intellectual disability, reinforcing its relevance during neurodevelopment[16]. Genetic factors are estimated to account for approximately 64–91% of ASD heritability[17,18] underscoring the importance of identifying and functionally characterizing ASD-associated genes, such as WAC. Although WAC mutations cause DESSH and are strongly linked to ASD, the molecular mechanisms by which WAC dysfunction leads to neurodevelopmental phenotypes remain poorly defined.

In addition, emerging evidence implicates widespread dysregulation of multiple neurotransmitter systems in ASD[19], including the cholinergic system and monoaminergic pathways involving dopamine (DA)[20] and serotonin (5-hydroxytryptamine; 5-HT)[20,21]. Alterations in acetylcholine (ACh) signaling, including changes in acetylcholinesterase (AChE) activity and acetylcholine receptor (AChR) expression, have been reported in ASD and linked to deficits in attention, cognition, and sensory processing[22,23].

Postmortem studies further support cholinergic system dysfunction, revealing reduced expression of nicotinic acetylcholine receptors (nAChRs) in the frontal and parietal cortices of individuals with ASD[20,24-26]. Among nAChR subtypes, the *α*7 nicotinic receptor (CHRNA7) has received particular attention due to its established roles in sensory processing, cognition, working memory, and attention, as well as its high expression in brain regions critical for these functions, including the hippocampus and frontal cortex[27,28] and its association with ASD[29,30]. In addition, reduced levels of choline, the precursor of acetylcholine and an agonist for nicotinic acetylcholine receptors (nAChR), have been observed in ASD patients[31], with symptom severity correlating with decreased cytosolic ACh levels[32]. Consistent with findings in individuals with ASD, studies using ASD-relevant animal models further demonstrate the relevance of cholinergic dysfunction to behavioral phenotypes associated with ASD.

On the other hand, serotonin signaling is notably complex in ASD. While a substantial subset of individuals exhibits hyperserotonemia in peripheral blood, central serotonergic signaling in the brain appears reduced or developmentally altered, indicating compartment-specific dysregulation rather than a uniform elevation [21,33]. Elevated blood serotonin levels have been reported in autism patients and in animal models compared with controls[21,34] In contrast, postmortem studies provide evidence for reduced binding of 5-HT2A and 5-HT1A receptors in ASD brain tissue[21,34], suggesting a potential feedback response to curtailed serotonin signaling. Together with reports of altered 5-HT synthesis, transporter function, and receptor expression, these findings suggest that disrupted serotonergic signaling during early brain development may contribute to atypical cortical circuit formation, sensory processing deficits, and social behavioral impairments.

Dopaminergic dysfunction has also been implicated in ASD, particularly in pathways governing reward processing, motivation, executive function, and repetitive behaviors[20,35-37]. Several studies report altered DA receptor expression, dopamine transporter activity, or mesolimbic circuit function in ASD[19,37,38]. However, these findings are heterogeneous and, in some cases, contradictory, reflecting differences in developmental stage, brain region, and methodological approach. Collectively, these observations underscore that DA and 5-HT signaling abnormalities in ASD are highly context-dependent and developmentally regulated, reinforcing the need to consider neuro-modulatory imbalance as a contributing rather than singular mechanism underlying ASD pathophysiology. These complexities will be further addressed in the discussion section in relation to genetic and molecular perturbations affecting neurodevelopment. Given that WAC regulates genes involved in neuronal signaling, it is plausible that WAC dysfunction disrupts multiple neurotransmitter systems, culminating in a complex neuro-behavioral phenotype. Significant knowledge gaps exist with respect to the relationship between WAC and neurotransmission. The nematode *Caenorhabditis elegans* offers a genetically tractable model for investigating the functional consequences of WAC ortholog disruption, given its conserved neurotransmitter pathway and the presence of WAC-like genes, *wac-1.1* and *wac-1.2* (e.g., PHX2587 [*wac-1.1*&*wac-1.2(syb2587)*]). In *C. elegans*, cholinergic neurotransmission mediated by acetylcholine receptors (AChRs) is critical for locomotion and neuromuscular signaling and is extensively interconnected with other neuro-modulatory systems, including DA and 5-HT, to regulate coordinated behavioral output. These processes can be quantitatively assessed through established behavioral paradigms to observe cholinergic synaptic function[39,40] as well as broader circuit-level neurotransmitter interactions. This study aimed to characterize the role of *wac* loss-of-function in modulating neurobehavioral outcomes to elucidate how *wac* dysregulation may contribute to altered neural signaling.

## Material and Methods

### Culture and maintenance of Strains

*Caenorhabditis elegans* strains, N2 (wild type) and PHX2587 (*wac-1.1*&*wac-1.2*(syb2587)) and *Escherichia coli* OP50, were procured from the Caenorhabditis Genetics Centre (University of Minnesota, Minnesota), grown on Nematode growth medium (NGM), and cultured at 22°C. A synchronized population of worms was obtained by sodium hypochlorite treatment. Embryos were incubated overnight at 15°C in M9 buffer to obtain L1 worms. RNAi experiments were conducted as described by Ahringer 2006[41]. Briefly, RNAi clones were cultured in Luria broth supplemented with carbenicillin (25 µg/mL) and then seeded onto NGM-IPTG + Ampicillin plates (1 mM IPTG; 50 µg/mL Ampicillin), incubated overnight at 37°C. L1 worms were added to the plates the next day and allowed to grow for 48 hours at 22°C. HT115 with L4440 plasmid was used as an empty vector control (EV).

### Aldicarb assay for assessment of cholinergic transmission

Determination of cholinergic transmission was conducted as described[40]. Briefly, adult N2 and PHX2587 worms were washed 3 times and transferred to NGM-Aldicarb plates (0.5 mM Aldicarb). Aldicarb, an acetylcholinesterase inhibitor, causes buildup of ACh, resulting in the flexion of muscles. The percentage of worms paralyzed at a given time is indicative of ACh signaling. The number of paralyzed worms was counted every 30 minutes, and the percentage of paralyzed worms was calculated. Any worms that were lost or injured were excluded. The percentage of paralyzed worms was compared to the wild type worms, N2. The Aldicarb assay has been utilized to study functional alterations resulting from perturbed synaptic transmission[42]. Previous studies have shown its ability in the assessment of cholinergic transmission in phytomolecules and established pharmacological AChE inhibitors such as donepezil[39,43].

### Assay for nicotinic acetylcholine receptor activity

Levamisole assay was used in combination with Aldicarb assay, to assess the activity of nicotinic acetylcholine receptor (nAChR) as described previously[39,40,44] with slight modifications. Levamisole is an allosteric modulator of nAChR[45], which induces paralysis in nematodes[46]. Elevated nAChR activity is evident from increased paralysis at a given time point[39,40]. Briefly, the age-synchronized adult N2 and PHX2587 worms were washed from the NGM-OP50 plates. Approximately 40 worms were added to the wells of 96-well plates. An equal volume of 500 µM Tetramisole HCl (Levamisole) solution (dissolved in M9 buffer) was added, and worms were analyzed for paralysis every 10 minutes. The percentage of the worms paralyzed was calculated for the time point when approximately 50% of the worms were paralyzed in N2 worms.

### Assay for motility

Motility assessment was conducted using thrashing assay as described previously[47] with slight modifications. Briefly, single worms were transferred to each well of 96-well plate in 50 µL M9 buffer and allowed to acclimatize for 10 minutes. The number of thrashes was scored for a period of 30 seconds. A head-to-tail sinusoidal movement was counted as a single thrash. A total of 20 worms were employed in each biological replicate, with an interval of 10 minutes after every 5 worms were transferred. The average number of thrashes was calculated for 20 worms and was used to perform statistical analysis.

### Relative quantification of ACh level and AChE activity

ACh levels and AChE activity were determined using the Amplex Red Acetylcholine/Acetylcholinesterase Assay kit, following the manufacturer’s protocol (Thermo Fisher Scientific, Cat. No.: A12217). The estimation of ACh levels was performed as previously described[39]. Briefly, age-synchronized adult, N2 and PHX2587 worms were washed and sonicated in a 1× reaction buffer. The worm suspension was then centrifuged at 7000 rpm for 7 minutes. One hundred microliters of supernatant were added to 100 µL of the reaction mixture (prepared by adding 200 µL of 20 mM Amplex Red solution, 100 µL of 200 U/mL HRP solution, 100 µL of 100 U/mL acetylcholinesterase solution, and 100 µL of 20 U/mL choline oxidase solution in 10 mL q.s. of 1× reaction buffer) in black well plates. After 30 minutes of incubation at room temperature, fluorescence was measured using a fluorimeter (BioTek Synergy H1 microplate reader, St. Louis, MO) with excitation and emission at 544 and 590 nm, respectively.

For AChE activity estimation, 100 µL of the supernatant was added to 100 µL of the reaction mixture consisting of 200 µL of 20 mM Amplex Red solution, 100 µL of 200 U/mL horseradish peroxidase solution, 10 µL of 100 mM ACh solution, and 100 mL of 20 U/mL choline oxidase solution in 10 mL q.s. of 1× reaction buffer. The relative fluorescence for ACh levels and AChE activity (measured at excitation: 545 nm and emission: 590 nm) was normalized against protein content, calculated using the BCA assay[48].

### 1-nonanol assay

Dopamine-associated behavior was evaluated using 1-nonanol-based repulsive behavior[49]. Repulsive behavior towards odorants has been reported in the literature[40,50-53]. Worms with optimum dopamine levels exhibit repulsive behavior towards 1-nonanol, while worms with curtailed dopamine show prolonged time to avert olfactory stimulus. A decreased repulsion time is indicative of higher dopamine levels and vice versa[53,54]. The assay was performed as previously described[40,53]. Briefly, the adult worms were washed 3 times with M9 buffer and placed on NGM plates. The poking lash dipped in 1-nonanol was positioned close to the head region of the worms, while ensuring not to touch the worms avoiding a mechanosensory response. Any worms prodded accidentally or exposed to excess 1-nonanol were excluded to rule out interference. Repulsion time, i.e. time taken for the worms to exhibit repulsive behavior was calculated using a stopwatch. The stopwatch was started at the time of the exposure and stopped when the worm reversed and turned its head 45 degrees.

### RNA isolation, cDNA synthesis, and quantitative real time PCR

Total RNA was extracted from N2 and PHX2587 worms using RNAzol reagent (Molecular Research Centre Inc.) as per manufacturer’s instructions. Briefly, worms were washed with nuclease-free water and homogenized in 200 µL of RNAzol on ice. The homogenate was incubated at room temperature for 10 minutes, and centrifuged at 13,000 rpm for 10 minutes, 4°C. Supernatant was mixed with 100 µL of nuclease-free water. After vortexing for 15 seconds, tubes were incubated at room temperature for 15 minutes. RNA was then precipitated by adding an equal volume of isopropyl alcohol and incubation for an additional 15 minutes at room temperature. Finally, RNA was pelleted by centrifugation at 13,000 rpm for 10 minutes at 4°C. The RNA pellet was washed two times using 75% chilled ethanol by centrifugation at 6,500 rpm for 5 minutes at 4°C. RNA quantity and quality were assessed using a nanodrop. cDNA was synthesized using 1 µg of RNA in a thermal cycler using the RevertAid H Minus First Strand cDNA synthesis Kit (Thermo Scientific) per manufacturer’s protocol. qRT-PCR was performed using the Bio-Rad CFX96 Real-Time PCR Detection System according to the manufacturer’s instructions. Differential expression was calculated using the 2^-ΔΔCT^ method[55]. Gene, g*pd-1* was used as internal control and used to calculate fold change. All primers were procured from Integrated DNA Technologies, with details as shown in Table S1 (orientation of oligonucleotides: 5’ to 3’).

### Animal model

The mutant mouse line used in this study has been previously described and characterized[56]. To generate constitutive *Wac* heterozygous mice, β-actin–Cre mice[57] were crossed with *Wac*^flox^ mice. Following germline recombination, wild-type (WT) and *Wac*^+/-^ offspring were obtained. After recombination, *Wac*^+/-^ mice were backcrossed and maintained on a CD-1 background for at least three generations prior to experimental use. Mice were provided to our laboratory for the collection of cortical brain tissue for experiments. All animal procedures were approved by the Institutional Animal Care and Use Committee (IACUC) at Michigan State University and conducted in accordance with institutional guidelines.

### Western blotting

Cortical tissue samples were collected from wild-type (WT) and *Wac*^+/-^ mice. The WT group consisted of nine mice (six males and four females), and the *Wac*^+/-^ group also consisted of nine mice (four males and four females). For each biological sample, two technical replicates were analyzed, with each replicate processed in an independent experimental run.

Total protein concentrations were determined using a BCA protein assay kit (23227, Thermo Fisher Scientific, USA). Equal amounts of protein were electrophoresed on Mini-PROTEAN Tetra Vertical Electrophoresis Cell (1658004, Bio-Rad Laboratories, Inc.) with 4–20% Mini-PROTEAN® TGX™ Precast Protein Gels (4561096, Bio-Rad Laboratories, Inc.) and transferred to Nitrocellulose (NC) blotting membrane (10600001, Cytiva, USA). Membranes were blocked in 5% (w/v) bovine serum albumin (BSA) (37520, Thermo Fisher Scientific, USA) in phosphate buffer saline (PBS) for 1 hour at room temperature. Membranes were incubated with primary antibodies following rabbit polyclonal anti-Nicotinic Acetylcholine Receptor alpha 7 (ab216485, Abcam, 1:1,000) and mouse monoclonal anti-GAPDH (AM4300, Invitrogen, 1:40,000) diluted in 1% (w/v) BSA in PBS-T for 3.5 hours at room temperature, then with the goat anti-rabbit (and goat anti-mouse (926-68070) fluorescence secondary antibody (1:20,000) diluted in PBS-T for 2 hours at room temperature. The membranes were visualized using Li-Cor Odyssey CLx and analyzed using Image Studio version 5.5 from LICORbio™.

### Statistical Analysis

Statistical analysis was performed using GraphPad PRISM, Version 10 (1992–2025 GraphPad Software, Inc., La Jolla, CA, USA). Each experiment was repeated at least three times. The *p*-values were calculated using an unpaired t-test. For all experiments, *p* < 0.05 was deemed statistically significant.

## Results

### wac mutant nematodes exhibit curtailed dopamine-associated behavior and impaired motility

Dopaminergic signaling has been extensively linked to social behavior[58,59], and has significantly highlighted its relevance to ASD[60-62]. To assess the impact of *wac* gene on dopamine-associated behavior, we employed 1-nonanol assay, an established indirect measure of dopaminergic signaling in *C. elegans*, where increased repulsion latency reflects reduced dopamine levels. PHX2587 (*wac-1.1*&*wac-1.2*(syb2587)) mutants exhibited a significant increase in repulsion time compared with N2 controls (PHX2587: 1.701 ± 0.048 vs. N2: 1.000 ± 0.028; *p* < 0.0001), indicating curtailed dopamine-associated behavior in *wac* mutant nematodes (**Figure 1 A, B**), suggesting impaired dopaminergic signaling. Locomotion in *C. elegans* is a complex process that is governed through coordination between neurons and neurotransmitters such as dopamine[63,64], serotonin[64], acetylcholine[65,66], and GABA[67,68]. ASD has also been linked to dysregulation of motor-related control in the gastrointestinal tract, with reports of altered motility and enteric nervous system deficits[69,70]. We tested the effect of *wac* gene mutation on motility using the thrashing assay. We observed that some of the worms exhibit immobilization upon transfer to M9 buffer, the percentage of immobilized worms was significantly higher in *wac* mutants (49.667% ± 6.009, *p =* 0.0074) compared with N2 (6.667% ± 4.410). The thrashing assay was performed only for actively swimming worms. Consistent with previous findings, *wac* mutants exhibited reduced motility (49.23 ± 1.341, *p* < 0.0001) compared with N2 (66.72 ± 1.087), as evidenced by a significantly lower number of thrashes in 30 seconds (**Figure 1 C, E; 1 D, F**). Findings from the thrashing assay revealed reduced motility and aberrations in the transition from crawling to swimming behavior in *wac* mutants.

**Figure 1.**
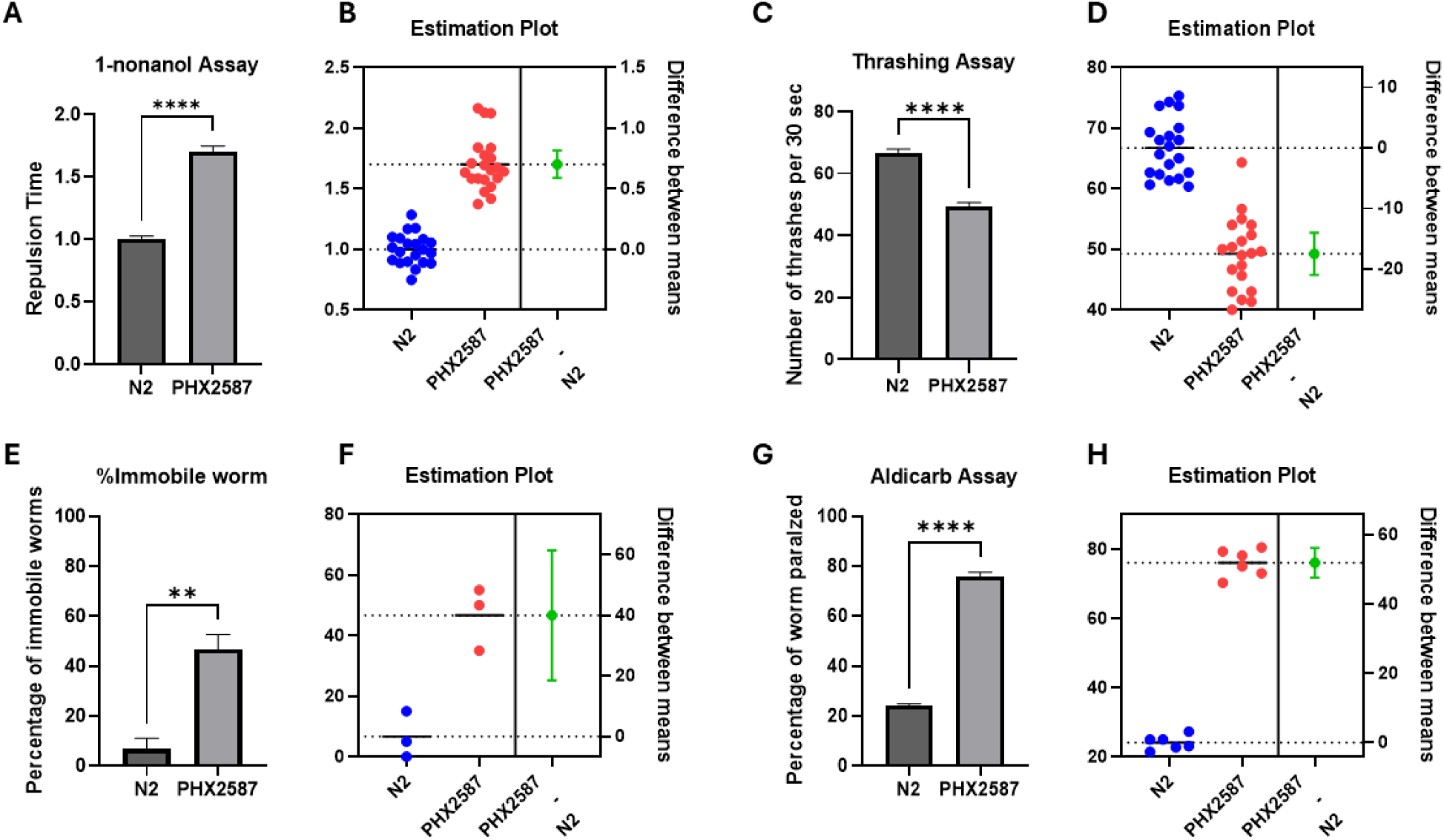
PHX2587 mutants exhibit impaired dopamine-dependent behavior, locomotor deficits, and increased cholinergic sensitivity. (A) Dopamine-dependent repulsion to 1-nonanol was significantly delayed in PHX2587 worms compared to wild-type N2, indicating reduced dopamine-mediated chemosensory behavior. Data represent mean ± SEM. ****p < 0.0001 (using Welch’s t-test) (B) Estimation plot of individual 1-nonanol repulsion times for N2 (blue) and PHX2587 (red). The right panel shows the mean difference between groups with a 95% confidence interval (green), confirming a significant increase in repulsion time in PHX2587. (C) PHX2587 worms displayed a significant reduction in thrashing frequency relative to N2, indicating impaired locomotor activity. Data are shown as mean ± SEM. ****p < 0.0001 (using Welch’s t-test) (D) Estimation plot of individual thrashing frequencies with the mean difference and 95% confidence interval (green), which does not cross zero, confirming decreased locomotion in PHX2587. (E) The percentage of immobile worms was significantly increased in PHX2587 compared to N2, further supporting locomotor impairment. Data represent mean ± SEM. **p < 0.01 (using Welch’s t-test). (F) Estimation plot of immobility showing individual values for N2 (blue) and PHX2587 (red). The mean difference with a 95% confidence interval (green) is shifted positively and does not cross zero, confirming an increased proportion of immobile worms in PHX2587. (G) PHX2587 mutants exhibited increased sensitivity to aldicarb-induced paralysis, shown as a higher percentage of paralyzed worms following aldicarb exposure compared to N2 controls, consistent with enhanced cholinergic neurotransmission. Data represent mean ± SEM. ****p < 0.0001 (Welch’s t-test). (H) Estimation plot of aldicarb-induced paralysis showing individual values and the mean difference with a 95% confidence interval (green). The confidence interval excludes zero, confirming increased aldicarb sensitivity in PHX2587.

### wac mutants exhibit enhanced cholinergic transmission, owing to heightened nicotinic acetylcholine activity

ASD is associated with abnormalities in the cholinergic system[24], as evidenced by altered expression of nAChR in the brains of affected individuals. To examine the impact of *wac* deletion on cholinergic neurotransmission, we first assessed acetylcholine-associated behavior using the aldicarb-induced paralysis assay. *wac* mutant worms exhibited significantly earlier paralysis compared with N2 controls (PHX2587: 76.028 ± 1.619 vs. N2: 24.084 ± 0.852; *p* < 0.0001), indicating hypersensitivity to aldicarb (**Figure 1 G, H**). This hypersensitivity to aldicarb suggests that *wac* deletion may dysregulate synaptic ACh release or turnover, consistent with elevated cholinergic activity. These results suggest that *wac-1.1* and *wac-1.2* deletion increases sensitivity to ACh accumulation, consistent with elevated synaptic ACh release or altered turnover.

Further, upon testing the effect on ACh levels and AChE activity, while a slight reduction in ACh levels and AChE activity was observed, it was statistically insignificant, negating the significant role of curtailed neurotransmitter and enzyme (**Figure 2 A, B**). Aldicarb assay informs the effect on total neurotransmission, whereas levamisole assay is used to ascertain the activity of nAChR[40,42]. In order to test the effect on nAChR activity we performed levamisole assay, and a significantly enhanced paralysis was observed in *wac* mutants, proving that the elevated cholinergic transmission was due to heightened nAChR activity only (**Figure 2 C, D**).

**Figure 2.**
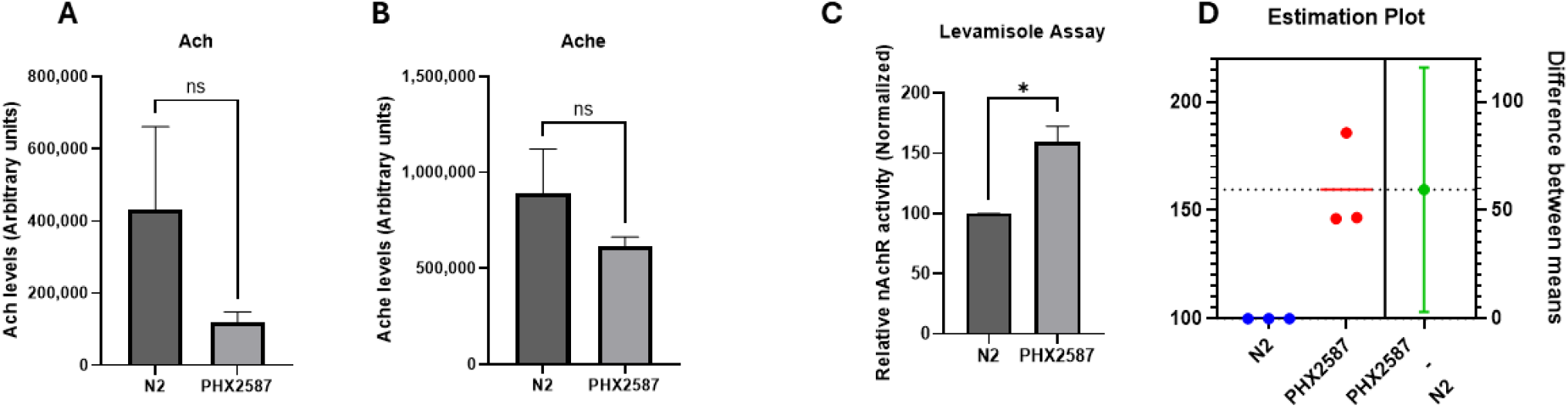
Altered cholinergic signaling in PHX2587 mutants without changes in total acetylcholine levels or acetylcholinesterase activity. (A) Quantification of acetylcholine (ACh) content in wild-type N2 and PHX2587 worms measured using the Amplex Red assay. PHX2587 exhibited a modest reduction in ACh levels relative to N2; however, the difference was not statistically significant. Data are presented in arbitrary units as mean ± SEM. (B) Acetylcholinesterase (AChE) enzymatic activity in N2 and PHX2587. A slight decrease in AChE activity was observed in PHX2587, but this difference did not reach statistical significance. Data are shown as mean ± SEM in arbitrary units. (C) PHX2587 mutants displayed significantly increased sensitivity to levamisole-induced paralysis compared to N2 controls. Relative paralysis responses, reflecting nicotinic acetylcholine receptor (nAChR) activity, were normalized and are presented as mean ± SEM. *p < 0.05 (unpaired t-test). (D) Estimation plot of levamisole-induced paralysis showing individual values for N2 (blue) and PHX2587 (red). The right panel presents the mean difference with a 95% confidence interval (green), which does not cross zero, confirming increased levamisole sensitivity in PHX2587.

Consistent with the aldicarb hypersensitivity observed in **Figure 1**, PHX2587 mutants displayed significantly greater levamisole-induced paralysis than wild-type N2. These results indicate that *wac-1.1* and *wac-1.2* deletion enhances nAChR-mediated responses, contributing to cholinergic hyperexcitability.

### wac mutants have significantly altered expression of genes involved in cholinergic transmission and reception

To test the effects on neurobehavioral aspects of cholinergic transmission and to identify the involvement of elevated nAChR activity, we investigated whether there were any alterations in genes related to cholinergic transmission. We quantified mRNA expression of 15 genes associated with acetylcholine synthesis, degradation, transport, and nicotinic acetylcholine receptor function, including *ace-1, ace-2, acr-2, acr-3, acr-12, cha-1, cho-1, lev-1, lev-8, lev-10, unc-17, unc-29, unc-38, unc-50*, and *unc-63*. A summary of gene expression changes is provided in **Figure 3**, with quantitative values detailed in **Table 1**.

**Figure 3.**
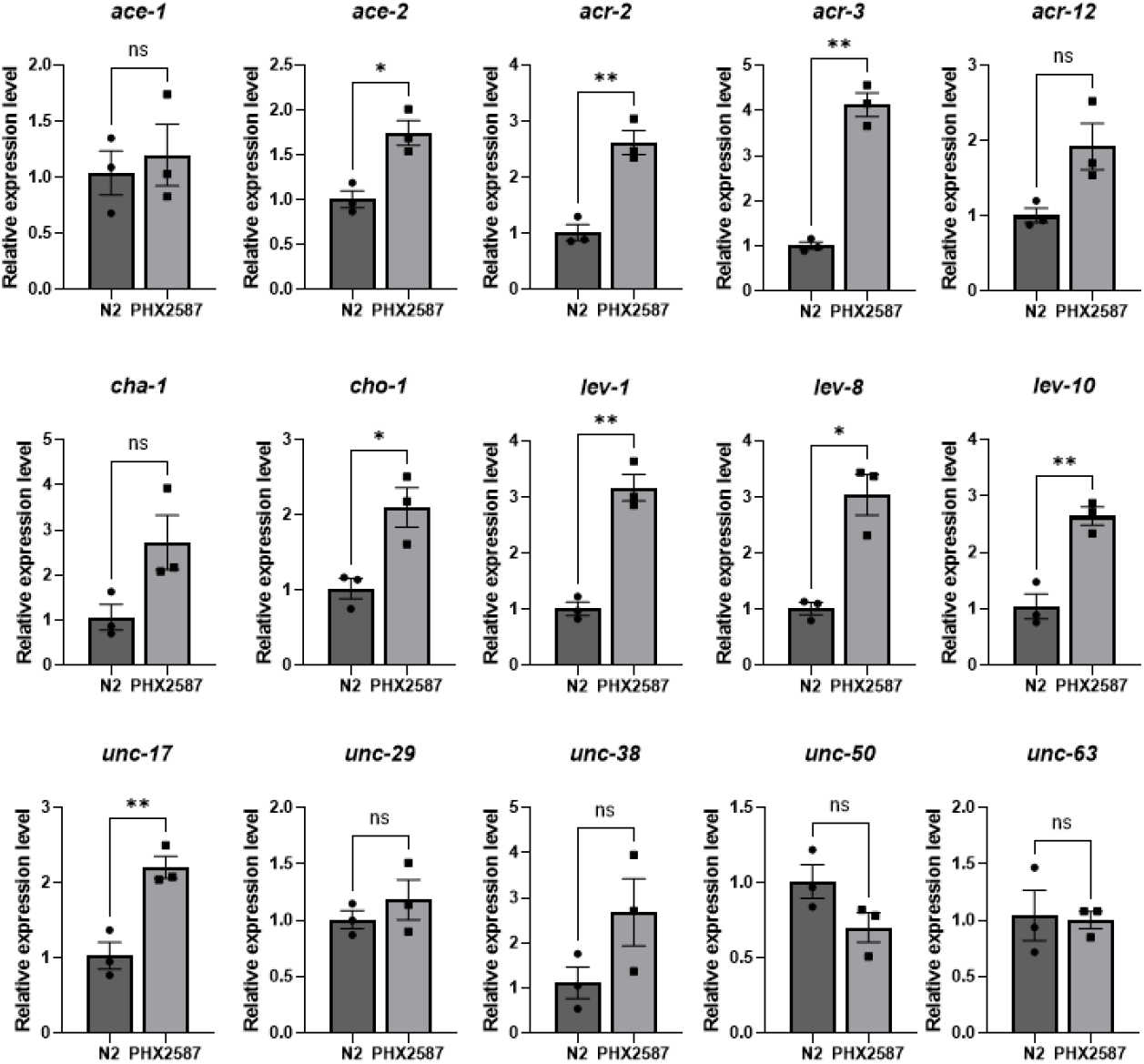
Quantitative PCR analysis of cholinergic gene expression in N2 and PHX2587 worms. Relative mRNA expression levels of cholinergic pathway genes were quantified by qPCR and normalized to N2 controls. Significant upregulation was detected in *ace-2, acr-2, acr-3, cho-1, lev-1, lev-8, lev-10*, and *unc-17*, while the remaining genes showed non-significant. Data represents mean ± SEM. Statistical significance was assessed using Welch’s t-test. *p < 0.05, **p < 0.01.

**Table 1.**
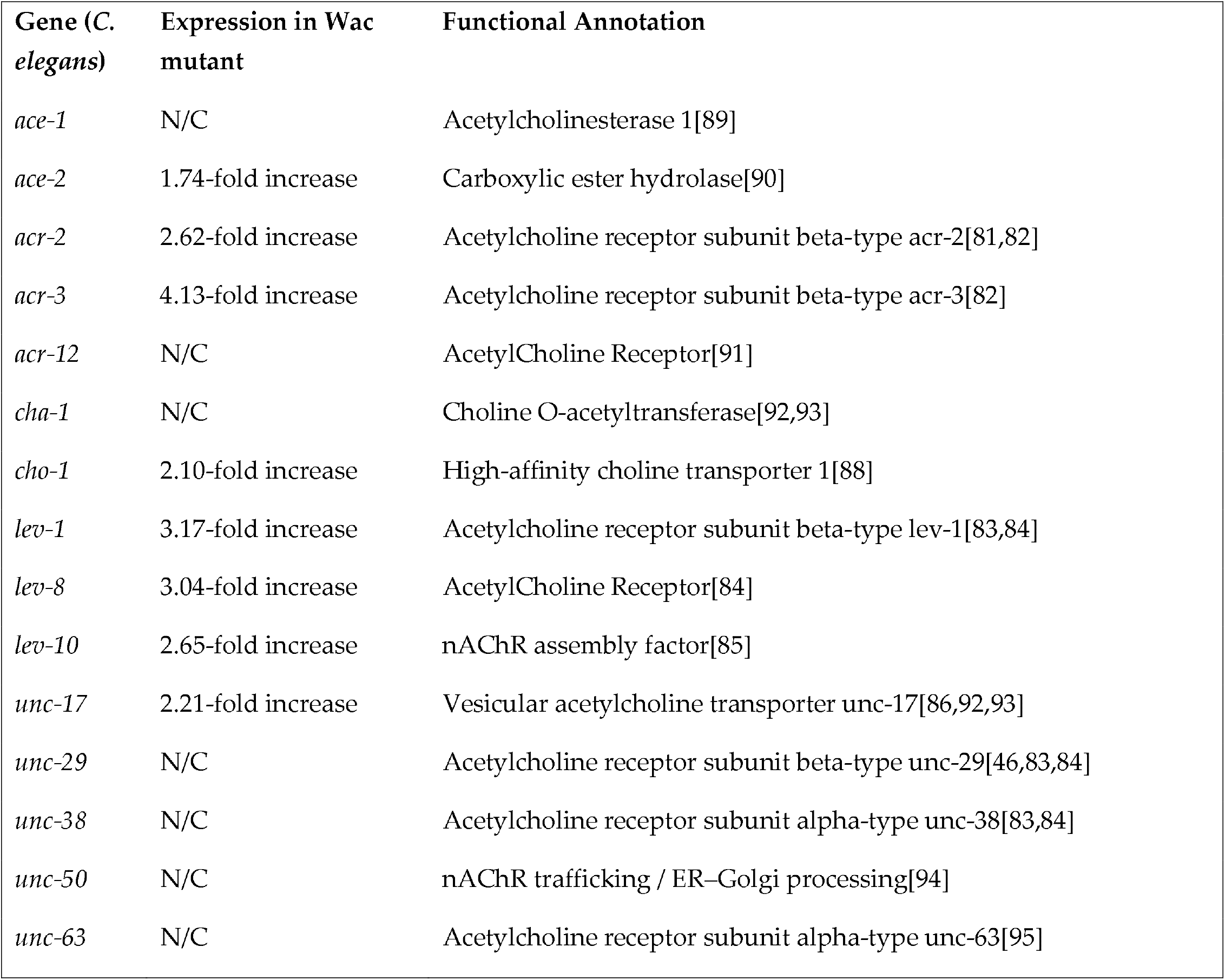
Cholinergic pathway genes and receptor-associated factors in C. elegans, their expression changes in the wac mutant PHX2587, and corresponding functional annotations. Genes involved in cholinergic neurotransmission or required for nicotinic acetylcholine receptor assembly and trafficking are listed along with their fold-change expression values in *PHX2587* relative to N2 controls. “N/C” indicates no significant change.

Notably, expression of *lev-1*, which encodes a non-α subunit of the levamisole-sensitive nAChR, was robustly upregulated, exhibiting a 3.17-fold increase relative to wild-type controls. Given the established role of *lev-1* in locomotion, egg-laying, and nAChR-mediated signaling, this upregulation provides a compelling molecular explanation for the enhanced sensitivity to levamisole and the elevated nAChR activity observed behaviorally. This suggests that *wac* deletion leads to receptor-level enhancement of cholinergic signaling, contributing to the hyperexcitability phenotype.

In PHX2587, multiple cholinergic genes, including *ace-2, acr-2, acr-3, cho-1, lev-1, lev-8, lev-10*, and *unc-17*, were simultaneously upregulated, suggesting that the transcriptional changes broadly affect components of the cholinergic signaling pathway.

### Lev-1 gene is not solely responsible for elevated nAChR activity

To determine whether *lev-1* upregulation contributes to enhanced cholinergic transmission in *wac*-deleted PHX2587 mutants, we performed *lev-1* RNAi in wild-type N2, and PHX2587, and performed aldicarb assay (**Figure 4)**. As expected, PHX2587 worms showed significantly increased sensitivity to aldicarb compared to control. Strikingly, *lev-1* knockout significantly decreased aldicarb sensitivity relative to control, consistent with its essential role in cholinergic transmission. Interestingly, silencing of *lev-1* in the PHX2587 background showed paralysis levels that were not significantly different from PHX2587. These results indicate that although *lev-1* is a critical component of nAChR, the *lev-1* silencing does not fully suppress the phenotype, suggesting that additional *wac*-dependent mechanisms are likely to contribute to the observed hypersensitivity.

**Figure 4.**
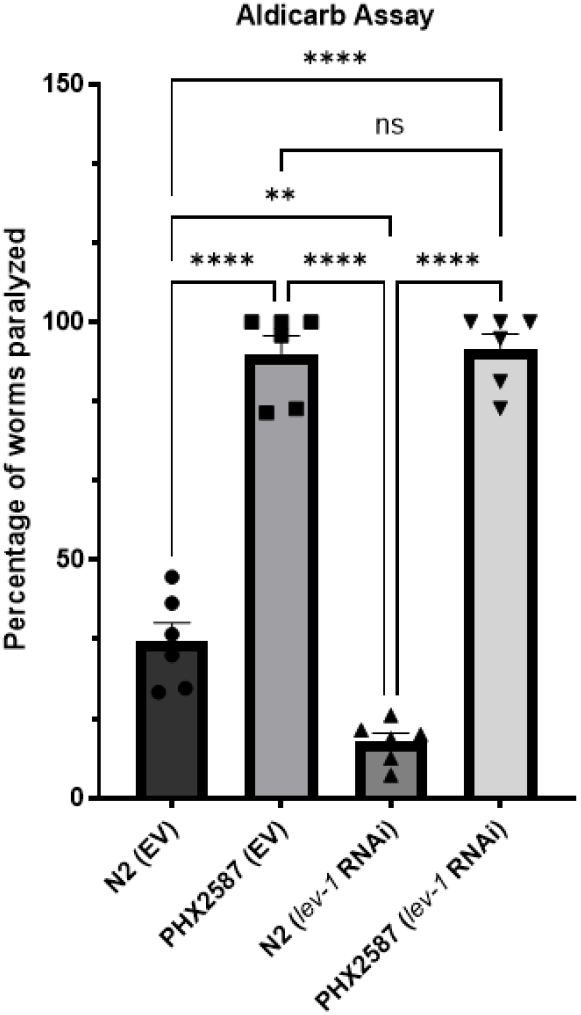
lev-1 is required for cholinergic hyperactivity in PHX2587 mutants. Aldicarb-induced paralysis was quantified in wild-type N2 and *PHX2587* worms carrying either an empty vector (EV) control or *lev-1* RNAi/knockout, as well as in *lev-1* single mutants and *lev-1*; *PHX2587* double mutants. *PHX2587* nematodes exhibited significantly increased aldicarb sensitivity compared to N2 controls, confirming cholinergic hyperexcitability. Loss of *lev-1* markedly reduced aldicarb sensitivity in both N2 and *PHX2587* backgrounds. However, aldicarb sensitivity in *lev-1*; *PHX2587* double mutants was comparable to *PHX2587* alone and remained significantly higher than *lev-1* single mutants. These results indicate that *lev-1* is required for the full expression of cholinergic hyperactivity in *PHX2587*, but *lev-1* deletion does not completely suppress the phenotype, suggesting the involvement of additional *wac*-dependent pathways. Data are presented as mean ± SEM and analyzed using one-way ANOVA followed by Dunnett’s post hoc test. **p<0.005, ****p < 0.0001.

### Wac Haploinsufficiency in Mice is associated with an increased CHRNA7/Wac ratio

To determine whether loss of murine *Wac* alters cholinergic receptor expression in vivo, we examined CHRNA7 protein levels in the cerebral cortex of *Wac*^+/-^ mice. Western blot analysis showed representative CHRNA7 immunoactivity bands in *Wac*^+/-^ mice compared with WT controls in the cortex (**Figure 5 A**). Quantification of CHRNA7 immunoreactivity normalized to GAPDH and expressed as fold change relative to WT did not reveal a statistically significant difference between groups; however, CHRNA7 levels in *Wac*^+/-^ mice showed a tendency toward increased expression relative to WT (**Figure 5 B**). To assess the extent of *Wac* haploinsufficiency, cortical *Wac* mRNA levels were quantified by qPCR. As expected, *Wac* mRNA expression was significantly reduced in *Wac*^+/-^ mice compared with WT controls (Figure 5 C). To account for inter-individual differences in *Wac* expression, we calculated an animal-wise ratio by dividing CHRNA7 protein fold change by *Wac* mRNA fold change (CHRNA7/*Wac*). This normalization revealed a significant increase in the CHRNA7/*Wac* ratio in *Wac*^+/-^ mice compared with WT controls (**Figure 5 D**). Surprisingly, some of the wild-type animals exhibited a lower expression, while on the other hand some of the *Wac*^+/-^ mice exhibited above 50% *Wac* expression. To make animal-wise normalization explicit, we tabulated CHRNA7 protein fold change, *Wac* mRNA fold change, and the derived CHRNA7/*Wac* ratio for each of the 18 mice (**Figure 5 E**). The same individual-level values are visualized in **Figure 5 F** to facilitate comparison across animals and to highlight the distribution of the CHRNA7/*Wac* metric.

**Figure 5.**
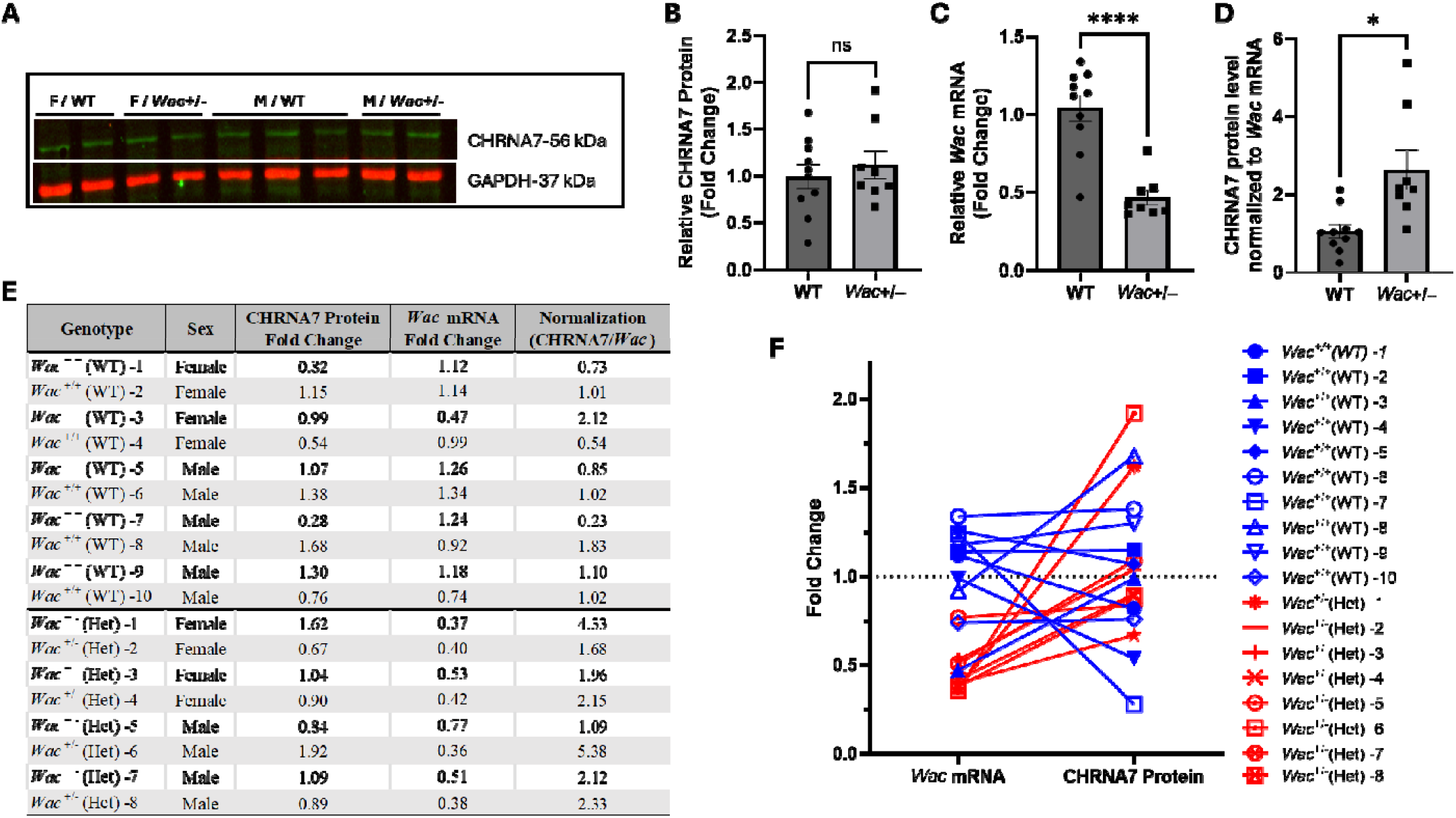
Wac haploinsufficiency is associated with an increased CHRNA7/Wac ratio in the mouse cerebral cortex. (A) Representative Western blot images showing CHRNA7 (upper panel, green) and GAPDH (lower panel, red) from cortical lysates of wild-type (WT) and *Wac*^+/–^ mice, separated by sex (female, F; male, M). Each lane represents an independent biological sample. (B) Quantification of CHRNA7 protein levels from Western blots. CHRNA7 intensities were normalized to GAPDH and expressed as fold change relative to WT controls. No significant difference was detected between genotypes. (C) qPCR validation of reduced *Wac* mRNA expression *Wac*^+/-^ mice displayed a significant reduction in *Wac* mRNA expression compared with WT controls. (D) Animal-wise normalization of CHRNA7 protein fold changes to *Wac* mRNA fold changes (CHRNA7/*Wac* ratio). This analysis revealed a significant increase in the CHRNA7/*Wac* ratio in *Wac*^+/–^ mice compared with WT controls. (E) Table summarizing individual animal values for CHRNA7 protein fold change, *Wac* mRNA fold change, and the derived CHRNA7/*Wac* ratio for all 18 mice. (F) Visualization of the same individual-level CHRNA7/*Wac* values shown in (E), illustrating the distribution across animals and facilitating comparison between genotypes. All data are presented as mean ± SEM. Statistical significance was determined using Welch’s t-test. *p < 0.05 and ****p < 0.0001.

## Discussion

ASD is a complex neurobehavioral and neurodevelopmental disorder[71], associated with dysregulation of several neurotransmitters, dopamine[63,64], serotonin[64], acetylcholine[65,66], and GABA[67,68]. The cause of ASD or its connection to a single genetic or environmental factor remains uncertain, although ASD is linked to both environmental and genetic factors. According to the SFARI Gene database (https://gene.sfari.org/), a literature-based resource for genes associated with autism spectrum disorder (ASD), the latest release (January 14, 2026) lists WAC as an ASD-related gene with a Gene Score of 1 and a syndromic classification. In this version of the database, 1,267 genes are listed as ASD-associated. Among them, 244 genes are assigned a Gene Score of 1, 312 genes are classified as syndromic, and 121 genes meet both criteria concurrently. In this study, using mouse and *C. elegans* models, we examined the effects of the *Wac* gene mutations on neurotransmitter imbalances associated with ASD, particularly acetylcholine and dopamine, with the aim of determining the correlation between the *Wac* gene and ASD.

Deficits in social development and communication are fundamental manifestations of ASD[38]. Accruing evidence identifies dysregulation of monoaminergic neurotransmitters, particularly dopamine and serotonin. Dopaminergic signaling has been associated with social behavior[58,59] and ASD[60-62]. Parallelly, serotonergic dysfunction has been associated with sensory processing, motor regulation, affective behaviors, and developmental abnormalities in ASD, underscoring the complex and context-dependent roles of these neuromodulators in the ASD brain[21,72]. Studies on dopamine-associated behavior using 1-nonanol assay showed reduced dopamine signaling, aligning with the symptoms in ASD. Locomotion in *C. elegans* is a complex process that is governed through coordination between neurons and neurotransmitters. Neurotransmitters such as dopamine[63,64], serotonin[64], acetylcholine[65,66], and GABA[67,68] play a critical role in nematode motility. Any alteration in neurotransmitter signaling is likely to exert motility defects. In ASD patients, impairment in motor-related control of the gastrointestinal tract [69,70] and in motor skills, balance, and gait [73-77] have been reported. Getting cues from the dopamine deficits, we tested whether there is any aberration in locomotion. We observed that *wac* mutants had reduced motility as evident through curtailed number of thrashes. We also observed a larger number of immobile worms in the case of *wac* mutants, indicating the inability of *wac* mutants to transition from crawling behavior to swimming behavior. More specifically, catecholamines, dopamine, and serotonin play a crucial role in the transition between crawling and swimming. Particularly, serotonin signaling is essential for crawl to swim transition and its maintenance[78]. This also suggests a possible reduction in serotonin levels. Conclusion from thrashing assay identified that reduced motility could be a confounding factor since the 1-nonanol assay is a movement-based behavior assay. To address this, we drew a comparison between the two assays; The reduction in motility was relatively lesser (26.21% only) while the increase in repulsion time was substantially higher (70.1%). This disparity indicates that while altered locomotion may contribute to delayed responses to a little extent, the pronounced impairment in the 1-nonanol assay primarily reflects disrupted dopaminergic sensory processing rather than generalized motor deficits.

ASD is also associated with abnormalities in the cholinergic system [25], as evidenced by altered nAChR expression in the brains of affected individuals. Postmortem analyses of autism patient brains have revealed region-specific changes in nAChR subtype expression, with marked reductions (approximately 40–50%) in α3, α4, and β2 nAChR subunits in the granule cell, Purkinje cell, and molecular layers of the cerebellum, while α7 nAChR expression was increased nearly threefold in the granule cell layer[79]. These findings suggest that, unlike disorders characterized by selective neuronal loss, ASD pathology may involve complex imbalances in receptor subtype distribution and signaling rather than uniform reductions in specific neuronal populations. Notably, *C. elegans* nAChRs share strong structural and functional similarity with vertebrate α7 nAChRs[80], supporting the relevance of this model for the study of cholinergic dysregulation in ASD. This prompted us to test the effect on ACh signaling. No effect on ACh levels and AChE activity was observed when assessed biochemically. One caveat of the biochemical assays is that they do not distinguish between synaptic and presynaptic ACh levels; behavioral assays are necessary to complement or validate findings about ACh signaling. Aldicarb assay indicates the effect on overall cholinergic neurotransmission through a paralysis-based phenotype, where a higher percentage of paralyzed worms suggests increased ACh signaling. Using the Aldicarb assay, we found that *wac* deletion resulted in enhanced cholinergic transmission. In order to check the involvement of nAChR, we performed levamisole assay that informs the activity of nAChR through a paralysis-based phenotype[40,42]. We observed enhanced levamisole-induced paralysis, indicating heightened nAChR activity. These findings suggest that the enhanced paralysis observed in PHX2587 is likely to result from increased postsynaptic sensitivity through upregulation of nAChRs. To further delve into the molecular mechanisms, we assessed the effect of *wac* gene deletion on the expression of genes responsible for Ach synthesis, transmission and reception. We quantified mRNA expression of 15 genes, *ace-1, ace-2, acr-2, acr-3, acr-12, cha-1, cho-1, lev-1, lev-8, lev-10, unc-17, unc-29, unc-38, unc-50*, and *unc-63* (**Figure 3, Table 1**). While the pro-signaling genes were upregulated, a similar countering response was observed among genes that can curtail cholinergic transmission. Upregulation in genes, *acr-2*(encoding non-alpha subunit of nAChR[81]), *acr-3* (encoding non-alpha subunit of nAChR[82]), *lev-1*(encoding non-alpha subunit of nAChR[83]), *lev-8*(encoding alpha subunit of nAChR[84]), and *lev-10*(encoding transmembrane protein localized to cholinergic neuromuscular junction[85]) can be connected to the elevated nAChR activity in addition to enhanced *unc-17* (encoding vesicular ACh transporter[86]) expression. A feedback response by the upregulation of *ace-2* (encoding Ache, responsible for degradation of ACh[87]), and *cho-1* (encoding choline transporter responsible for transport of ACh back to pre-synapse[88]) was also observed. Considering the heightened levamisole-induced paralysis, we tested the hypothesis concerning nAChRs by silencing one of the key nAChR related gene. The gene *lev-1* is known to produce the strongest resistance to levamisole amongst the *lev* gene family[83], hence we chose to test the effect of *lev-1* RNAi on aldicarb-induced paralysis in wild-type and *wac* mutant worms. While *lev-1* silencing did reduce aldicarb-induced paralysis, the percentage reduction in wild-type and *wac* mutant worms was insignificantly different, indicating the role of other key factors in enhanced cholinergic signaling.

To further validate and complement our findings, we measured CHRNA7 levels in mutant *Wac*^+/-^ mice. Initial results showed a trend of enhanced CHRNA7 expression in *Wac*^+/-^ mice, however the results were statistically insignificant and hence inconclusive. We further delved into it and performed qPCR to check the *Wac* mRNA expression in both wild-type and *Wac*^+/-^ mice. To our surprise, we noticed that some of the *Wac*^+/-^ mice exhibited less than 50% reduction in *Wac* expression, and some wild-type mice exhibited reduced *Wac* expression. On the one hand, we postulated that loss of one allele could possibly be eliciting genetic compensation[76,77], which could be due to the essential nature of the *Wac* gene and is further supported by the fact that homozygous *Wac* deletion leads to embryonic lethality[56]. Another possibly is that the WAC protein or *Wac* transcript could be dynamically regulated; some evidence for this comes from a study that found WAC protein levels were elevated by cyclin-dependent kinase 1[5]. Furthermore, we noticed inverse relationships between *Wac* expression and CHRNA7 levels in both the wild type and *Wac*^+/-^ mice. This pattern suggests that *Wac* dosage influences the relationship between CHRNA7 protein expression and baseline cellular context, pointing to a regulatory role for *Wac* in constraining CHRNA7 protein output. Presenting matched mRNA and protein measurements on a per-animal basis allows this relationship to be evaluated without assuming proportionality between transcript and protein levels. Together, these findings support a model in which reduced *Wac* expression is associated with relative upregulation of CHRNA7 protein, consistent with dosage-sensitive regulation of cholinergic signaling components. Together, these data demonstrate that partial loss of *Wac* leads to increased cortical CHRNA7 expression, supporting the idea that WAC acts as a negative regulator of nicotinic acetylcholine receptor (nAChR) subunits in the mammalian brain. This pattern mirrors our findings in *C. elegans*, where deletion of *Wac* resulted in up-modulation of the nAChR activity and enhanced cholinergic signaling, suggesting that *Wac*-repression, emanating enhanced cholinergic signaling, is evolutionarily conserved.

### Conclusion

Our studies elucidate the effect of the WAC gene deletion on neurobehavioral outcomes and evaluate the said gene for its potential relevance to the manifestations observed in ASD-associated neurobehavioral aberrations, mainly being up-modulation and down-modulation of cholinergic and dopaminergic signaling. Nematode models have emerged as an alternative model for testing the mechanisms involved. Future studies will examine in greater detail the mechanisms and stage-specific gene expression in molting related to cholinergic signaling and socio-behavioral assays concerning ASD.

## Acknowledgement

Funding for Dariangelly Pacheco-Cruz: DP-C was funded by the by the Pharmacological Sciences Training Program at Michigan State University, 5T32GM142521-04. A special mention of Mr. Veer Patel, who has always been an inspiration for me (SRS) to conduct research in this field. Strains were provided by the CGC, which is funded by NIH Office of Research Infrastructure Programs (P40 OD010440).

