## Supplementary data for "Neurobehavioral impacts of the autism risk gene, WAC: Studies involving C. elegans and Mice"

***Table S1: Sequence of the primers used***

|  | **Gene Name** | **Primer Sequence 5’ to 3’** |
| --- | --- | --- |
| **1** | ***gpd-1* F** | AAAGTCATTCCGGAGCTGAAC GGA |
|  | ***gpd-1* R** | AGCGGCCTTGACTACCTTCTTGAT |
| **2** | ***cha-1* F** | GGGAAAGGGAAGAAACGA |
|  | ***cha-1* R** | GGACCACTGCACCATAC |
| **3** | ***unc-17* F** | GGCTCCCACCATTCTTTC |
|  | ***unc-17* R** | GCATCACCGATGTGGTATAG |
| **4** | ***ace-1 F*** | CAGAGTGAGGACACTTACTTTGGA |
|  | ***ace-1 R*** | CCCAAACCATTACAGCCAATTTCT |
| **5** | ***ace-2* F** | GCCCATTCGGATTTCTCTAC |
|  | ***ace-2* R** | AGCTGATTCTCCGAACAAAG |
| **6** | ***cho-1* F** | TCGATTCCACCGGATAAGA |
|  | ***cho-1* R** | CAAACATTGATGCTGCTGATAG |
| **7** | ***acr-2* F** | CAGGAATATGGGACGTGATTG |
|  | ***acr-2* R** | GAGAACCGTTGGGATGATAAG |
| **8** | ***acr-3* F** | TAATAGATGCACCGGGTTTG |
|  | ***acr-3* R** | ACTGGATTCTGCTGGTAAATA |
| **9** | ***acr-12* F** | ACGGGTAGATATGTGGATTTG |
|  | ***acr-12* R** | CGTTCGGATGTCAAGGATAG |
| **10** | ***lev-1* F** | CGCAGAGACGAAGAGATTAC |
|  | ***lev-1* R** | CTGATGGAGCAGTGAAGATTAT |
| **11** | ***lev-8* F** | GTGGATACCACAACGGATAAG |
|  | ***lev-8* R** | GAACTGGTCTGACAGCTTTAT |
| **12** | ***lev-10* F** | ACGACACATCCAGCAAAG |
|  | ***lev-10* R** | GTCTCTCGATTGCTCTCAAC |
| **13** | ***unc-29* F** | CGAGGACCAAGAACTCATC |
|  | ***unc-29* R** | ACTGTCCAACTCCTGGTA |
| **14** | ***unc-38* F** | CGCTGACAGCAACTACA |
|  | ***unc-38* R** | CAGGAGCCGAACTTCAA |
| **15** | ***unc-50* F** | CATCCCAGTCACCGATTT |
|  | ***unc-50* R** | TGAGAGCTTGGCGAATG |
| **16** | ***unc-63* F** | GGTCATCATAGCAAACCAAATC |
|  | ***unc-63* R** | GAGTTGTCGCGTGGTAAA |
| **17** | ***Wac F*** | CAGATGATTGGTCTGAGCACATTAG |
|  | ***Wac R*** | TTTCAAGCCACTCTTTTGGTTTCT |
